# Environmental correlates of the European common toad hybrid zone

**DOI:** 10.1101/806067

**Authors:** Jan W. Arntzen, Daniele Canestrelli, Iñigo Martínez-Solano

## Abstract

The interplay between intrinsic (development, physiology, behavior) and extrinsic (landscape features, climate) factors determines the outcome of admixture processes in hybrid zones, in a continuum from complete genetic merger to full reproductive isolation. Here we assess the role of environmental correlates in shaping admixture patterns in the long hybrid zone formed by the toads *Bufo bufo* and *B. spinosus* in western Europe. We used species-specific diagnostic SNP markers to genotype 6584 individuals from 514 localities to describe the contact zone and tested for association with topographic, bioclimatic and land use variables. Variables related to temperature and precipitation contributed to accurately predict the distribution of pure populations of each species, but the models did not perform well in areas where genetically admixed populations occur. A sliding window approach proved useful to identify different sets of variables that are important in different sections of this long and heterogeneous hybrid zone, and offers good potential to predict the fate of moving contact zones in global change scenarios.

## Introduction

The study of hybrid zones, or areas where individuals from two genetically differentiated population lineages meet and produce hybrid offspring, is critical to our understanding of how new species form and remain distinct through evolutionary time (Barton & Hewitt, 1985; Harrison & Larson, 2014). The outcome of this admixture process ranges from complete genetic merger to full reproductive isolation with strong selection against hybrid individuals, and is a result of the interaction between intrinsic and extrinsic factors (Mallet, 2005; Abbott et al., 2013). Among the former are developmental, physiological and behavioral traits contributing to reproductive isolation, whereas extrinsic factors include those associated with the spatial and climatic context of the hybrid zone.

Hybrid zones often extend for hundreds of kilometers across heterogeneous landscapes, with features like mountains, rivers, soil types or roads differentially restricting dispersal and, consequentially, the degree of admixture in alternate sections of the contact zone. Similarly, spatially varying climatic conditions along the hybrid zone potentially impose different selective pressures on parental species and their hybrids in different transects. While topography is generally more conserved through time, climatic changes operate at temporal scales that are relevant for the study of hybrid zone dynamics and have been invoked to explain patterns of asymmetrical introgression in moving hybrid zones. The study of the complex interplay of topographic and climatic factors varying over time and space in hybrid zones can thus illuminate the relative role of extrinsic factors in reproductive isolation, explain discordant genetic patterns in different sections of a hybrid zone, and help predict the fate of hybrid zones (i.e., the stability of reproductive isolation through time) in the face of climatic changes (McQuillan & Rice, 2015; Taylor et al., 2015; Hunter et al., 2017; Ryan et al., 2018).

The European common toad hybrid zone is an emerging model system in the study of speciation. The common (*Bufo bufo*) and spined (*Bufo spinosus*) toads meet along a ca. 900 km hybrid zone from the Atlantic coast of northwest France to the Mediterranean coast of southeast France and northwest Italy (Recuero et al., 2012; Arntzen et al., 2018). This hybrid zone has been previously characterized with mitochondrial and nuclear DNA markers, which have revealed different patterns of genetic admixture across the hybrid zone, with transects in northwestern France resembling a classical tension zone model with selection against hybrids and those in the southeast of France and northwest of Italy characterized by strong cyto-nuclear discordance, probably as a result of spatial displacement of the hybrid zone in the past due to climatic changes (Arntzen et al., 2016, 2017; van Riemsdijk et al., 2019ab).

While previous studies have discussed the potential role of intrinsic factors in maintaining reproductive isolation across the hybrid zone, the relative role of environmental correlates has not been assessed so far, with the exception of certain topographic features reported to be associated with genetic transitions in different section of the contact zone (Arntzen et al., 2017, 2018). Here we use environmental data and a comprehensive genetic dataset to investigate the role of extrinsic factors in maintaining species borders through space and time.

## Materials and methods

A total of 6584 tissue samples was obtained from adult and larval toads from 514 localities across western Europe, with the emphasis on the broad region of species contact across France and Italy. Sampling was somewhat thin at the Atlantic coast and it was unsuccessful over the lower cols of the French Alps (col de Montgenèvre at 1854 m a.s.l., colle della Maddalena at 1996 m, col de Tende at 1870 m). All individuals were studied for the four species-diagnostic SNP nuclear markers <u>bdnf</u>, <u>pomc</u>, <u>rag1</u> and <u>rpl3</u> (Arntzen et al., 2016). Individuals with data missing for more than one marker were excluded. Sample size ranged from 1-358 with an average of 12.8. A total of 610 individuals was studied in duplicate. Results were not the same in two cases (0.08%) and involved the score of ‘22’ (i.e. homozygous for the spinosus-allele) versus the heterozygous ‘12’ condition. Both observations were subsequently treated as unavailable. The total of missing data was 2.0%. A subset of 2524 individuals from 185 localities was studied for another 27 diagnostic SNP markers (as in van Riemsdijk et al., 2019b; Arntzen, 2019) to determine the center of the two species contact zone more accurately. These latter localities were arranged in eight transects with hybrid zone centers located in France or Italy. For locality information, sample sizes and molecular species identifications see supplementary information I [not provided]. Mitochondrial DNA and toad morphology were not studied because for these characters species diagnostic performances break down over parts of the species’ parapatric range border (Arntzen et al., 2017, 2018).

The genetic data were subjected to Bayesian clustering with Structure software (Pritchard et al., 2000) with a predetermined K-value of two, corresponding to the number of species involved. The position of the mutual range border of *B. bufo* and *B. spinosus* was estimated by spatial interpolation of the Structure Q-values, to obtain the Q=0.5 isoline. In this procedure data were weighted by sample size and by the number of markers employed. We used Dirichlet cells with ILWIS (ILWIS, 2009) and linear interpolation with MyStat (Systat, 2007), which methods implied a weak and a strong spatial smoothing, respectively.

Environmental data considered include altitude and 19 climate variables from the WorldClim global climate database v2, available at http://www.worldclim.org (Hijmans et al., 2005; see also Fick & Hijmans, 2017). The parameter ‘slope’ was derived from altitude through a set of filter operations. Soil properties data are from the ESDA European soil Database v2.0, available at https://esdac.jrc.ec.europa.eu (see also Panagos et al., 2012) and were used as far as parameter values could be ordered. Vegetation and land use data were from the CORINE land cover database of the European Environment Agency, available at https://www.eea.europa.eu/publications/COR0-landcover. Data were grouped in the four classes ‘crop growing’, ‘forestation’, ‘pasture’ and ‘other’ (including e.g. wetlands, urban, industrial and the transport network).

To identify and subsequently reduce colinearity among the environmental variables we constructed the half-matrix of their pairwise Pearson correlation coefficients. This matrix was subjected to clustering with UPGMA. Variables were retained using criteria of partial independence at r<0.7 and selected in alphanumeric order (supplemental information II) [not provided]. The 20 variables considered for the construction of species distribution models are listed in table 1.

**Table 1.** Environmental variables considered for the construction of two-species distribution models for the common toad *B. bufo* and the spined toad *B. spinosus*.

| Class | Environmental variable (unit) | Code |
| --- | --- | --- |
| Climatic – WorldClim database |  |  |
|  | Annual mean temperature (C) | Bio01 |
|  | Mean diurnal range (C) | Bio02 |
|  | Isothermality | Bio03 |
|  | Temperature seasonality | Bio04 |
|  | Minimum temperature of coldest moth (C) | Bio06 |
|  | Mean temperature of wettest quarter (C) | Bio08 |
|  | Mean temperature of driest quarter (C) | Bio09 |
|  | Annual precipitation (mm) | Bio12 |
|  | Precipitation of driest month (mm) | Bio14 |
|  | Precipitation seasonality | Bio15 |
| Topographical – WorldClim database |  |  |
|  | Altitude (m a.s.l.) | Alt |
| Soil properties - ESDAC database |  |  |
|  | Dominant limitation to agricultural use<br>(recoded as no limitation, gravelly, stony and lithic) | Aglim1 |
|  | Total available water content (mm, 7 classes) | Awc |
|  | Bulk density (g cm<sup>-3</sup>, 10 classes) | Bd |
|  | Soil crusting (very weak to very strong, 5 classes) | Crusting |
|  | Depth to rock (cm, 4 classes) | Dr |
|  | Topsoil easily available water capacity (mm/m, 4 classes) | Eawctop |
|  | Sand (%) | Sand |
|  | Annual average soil water regime<br>(depth in cm and duration in months, 4 classes) | Wr |
| Land use and vegetation (categorical) – Corine database |  |  |
|  | Crop growing (classes 12-17, 19-21) | Crop |
|  | Forestation (classes 22-25) | Forest |
|  | Pasture (class 18) | Pasture |
|  | Other (classes 1-11,26-36, 40, 41) | Other |

We produced two-species distribution models, in which both species’ environmental attributes are contrasted under the assumption of a distribution in parapatry, as follows. First, we sampled environmental data for the localities of the genetically investigated toad populations within the area -5 to 12 degrees eastern longitude and 42 to 53 degrees northern latitude. Populations with Q<0.2 that represent *B. bufo* and Q>0.8 that represent *B. spinosus* were about equally frequent (*B. bufo* N=196, *B. spinosus* N=208). Data for 47 populations with intermediate Q-values were analyzed separately. Second, environmental data were sampled in a string of 17 overlapping and adjoining circular windows, positioned over the species contact zone, as defined by the Q=0.5 isoline (figure 1). A window diameter of 100 km was chosen to widely encompass the ca. 50 km wide *B. bufo - B. spinosus* hybrid zone (Arntzen et al., 2016, 2017, 2018; van Riemsdijk et al., 2019b). In consideration of scale, the string of windows followed the smooth curved species delineation (and not the more angular one; see below). The analyses involved 100 randomly chosen data points per window. The sampling localities were interpreted as pseudo-presences of *B. bufo* and *B. spinosus* if at the northeastern or at the southwestern side of the inferred species border, respectively.

**Figure 1.**
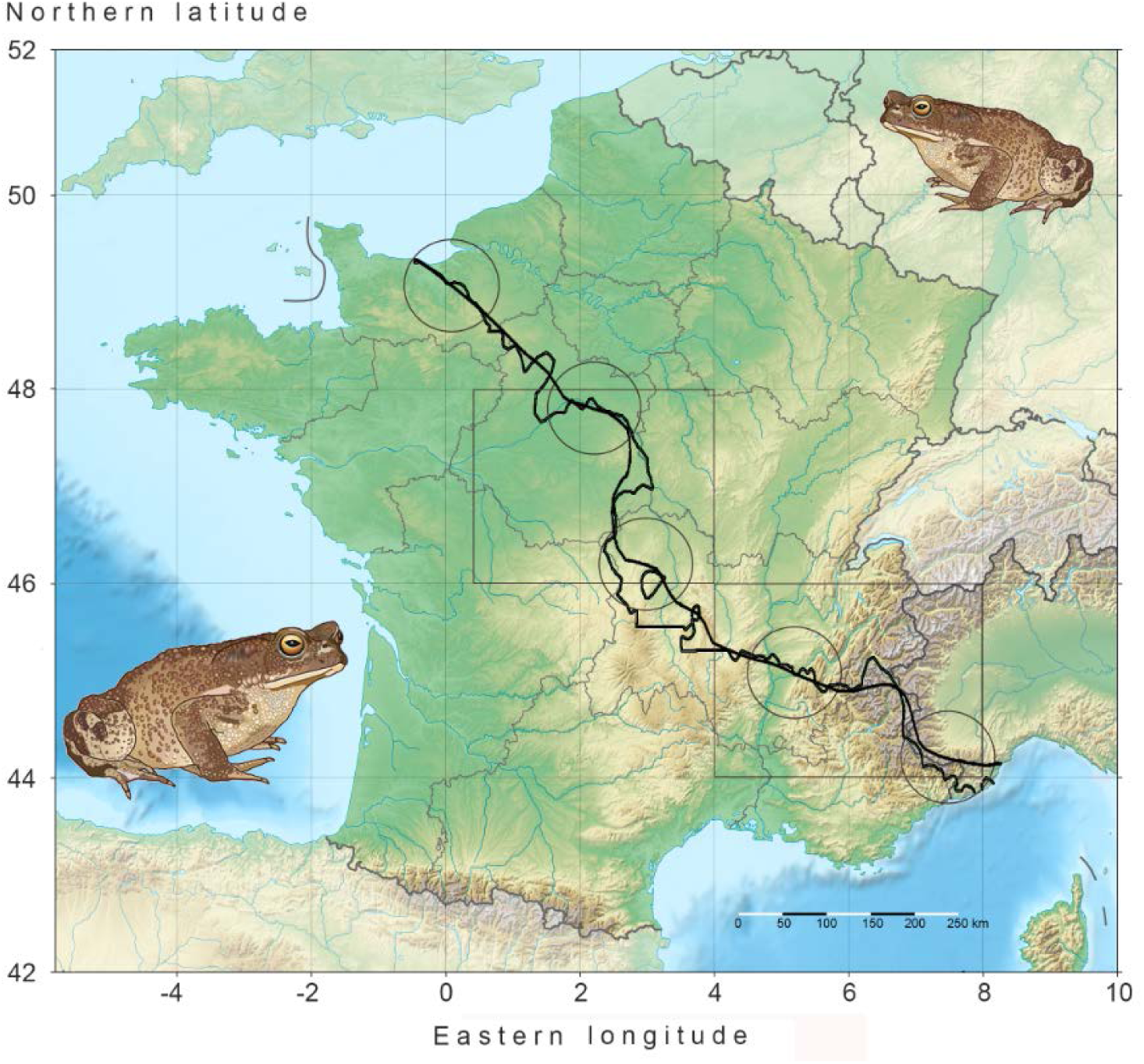
Western Europe with France and adjacent countries in Mercator projection. Colours from green to brown indicate increasing altitudes. The *Bufo bufo* versus *B. spinosus* mutual range delineation is based upon molecular genetic data, in which the smooth line is derived by linear interpolation whereas the more angular line is based upon Dirichlet cells (for details see text). The small bodied common toad *B. bufo* occurs to the northeast and the large bodied spined toad *B. spinosus* to the southwest of the mutual range border. Environmental data were gathered for 17 overlapping and adjoining windows positioned over the mutual range border. Here shown are window 1 in the northwest of France, window 17 in the northwest of Italy and windows 5, 9 and 13 in between. The two boxed areas are highlighted in figure 3. The base map was downloaded from MapsLand at https://www.mapsland.com, under a Creative Commons Attribution-ShareAlike 3.0 Licence. The animal drawings are by Bas Blankevoort, Naturalis Biodiversity Center.

Logistic regression analyses were performed with SPSS 20 (IBM SPSS, 2016) with ‘species’ as the dependent variable versus one categorical and 19 continuous environmental explanatory variables (table 1). Parameter selection was in the forward stepwise mode under the criteria of P_in_=0.05 and P_out_=0.10 under the likelihood ratio criterion. Distribution models were visualized with ILWIS (ILWIS, 2009). Model fit was assessed by the area under the curve (AUC) statistic.

## Results

The nuclear genetic data confirm that the hybrid zone of the common toad, *B. bufo* and the spined toad, *B. spinosus*, runs from the Atlantic coast to the Mediterranean (figure 1). The smooth and more angular estimates of the mutual species range delimitation run in parallel, with relatively large deviations from one another near the city of Orleans (window 4), the northern foothills of the Plateau Central (windows 9 and 10) and the French Alps (window 15). The selected two-species distribution model is represented by the logistic equation P_b_=(1/(1+exp(0.0167* alt+0.243* bio01+1.307* bio03+0.179* bio06-0.0357* bio08+0.0105* bio09-0.00856* bio12+0.0195* sand-69.899))), in which P_b_ is the probability of occurrence of *B.bufo* on a zero to unity scale (figure 2). The AUC model fit is 0.97. This model applied to 47 genetically admixed toad populations shows no significant correlation of the Structure Q-value and AUC model fit (r=0.157, P>0.05). Model fit over each of the 17 windows varied from ‘less than good’ (AUC<0.8) at windows 1, 6-8 and 14-16, to ‘good’ or ‘very good’ for windows 2, 3, 5, 12, 13 and 17 and approached unity at windows 4 and 9-11. Low model fit largely coincides with areas where i) sampling is thin (window 1, see supplementary information I) [not provided], ii) where rivers separate the species’ ranges, namely the Loire and the Cher at windows 5-8 (figure 3A) and the Rhône-Isère at windows 13-14, and iii) the Alps, where both species are thinly distributed (windows 15-16) (figure 3B). Evaluation of the performance of the eight selected variables over the remaining windows (i.e. windows 2-4, 9-12 and 17) shows qualitatively consistent results for altitude, bio03 and bio08, with lower values for *B. bufo* than for *B. spinosus* (figure 4). However, values vary markedly along the contact zone with, for example, the mutual species border being located at altitudes of ca. 200 m a.s.l. in the west of France, at ca. 500 m at the Plateau Central and at ca. 1200 m in the Ligurian Alps. The reverse situation with *B. bufo* at higher altitudes than *B. spinosus* was found at window 6, which coincides with the Loire river. The parameters bio09 and bio12 also show contrasting values over different sections of the contact zone. Finally, the parameters bio01, bio06 and sand show few or minor inconsistencies.

**Figure 2.**
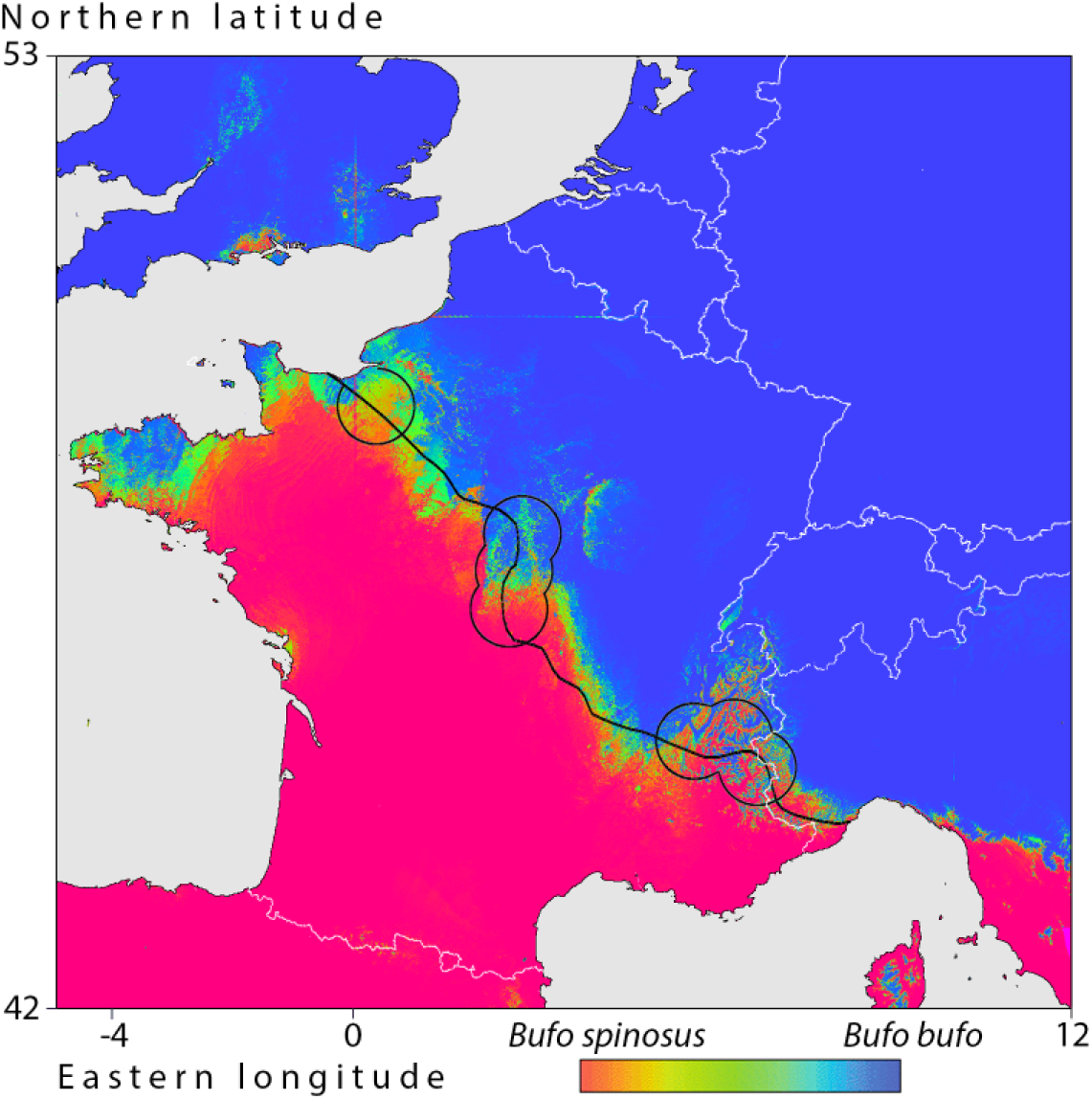
Two-species global distribution model for *Bufo* toads in western Europe. The colour scale runs from deep red for *B. spinosus* (P_b_ is zero) to deep blue for *B. bufo* (P_b_ at unity). The solid black line represents the center of the species’ hybrid zone from molecular data, as in figure 1. Outlined circular windows are those for which model fit is less than good (AUC<0.8, windows 1, 6-8 and 14-16).

**Figure 3.**
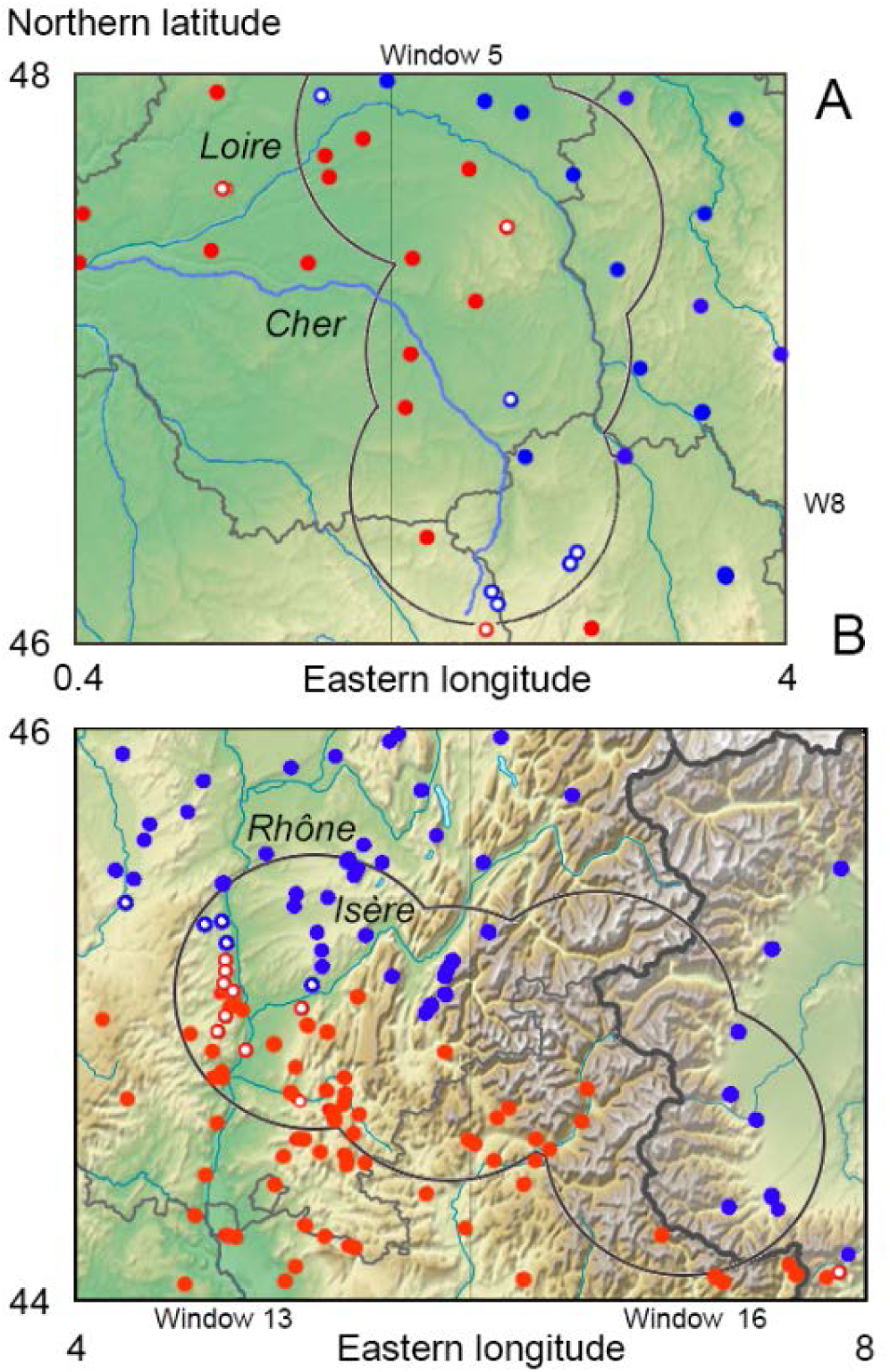
Two regions in the common-spined toad hybrid zone where the mutual species border appears to coincide with rivers. Coloured dots indicate toad populations with nuclear genetic species identifications as *Bufo bufo* (Q>0.5, blue symbols) and *B. spinosus* (Q<0.5, red symbols). Open dots have Q-values in the 0.2-0.8 range. For numerical detail see supplementary information I [not provided]. Base map figure credits as in figure 1. A) central France where the species border appears to coincide with the northernmost sections of the Loire (windows 5 and 6) and the upper stretches of the Cher (windows 7 and 8). B) southeastern France where the species border coincides with the Rhône and the lower Isère river at window 13. Note the paucity of data for the high Alps at windows 15 and 16 (see also Lescure and de Massary, 2012; Arntzen et al., 2017).

**Figure 4.**
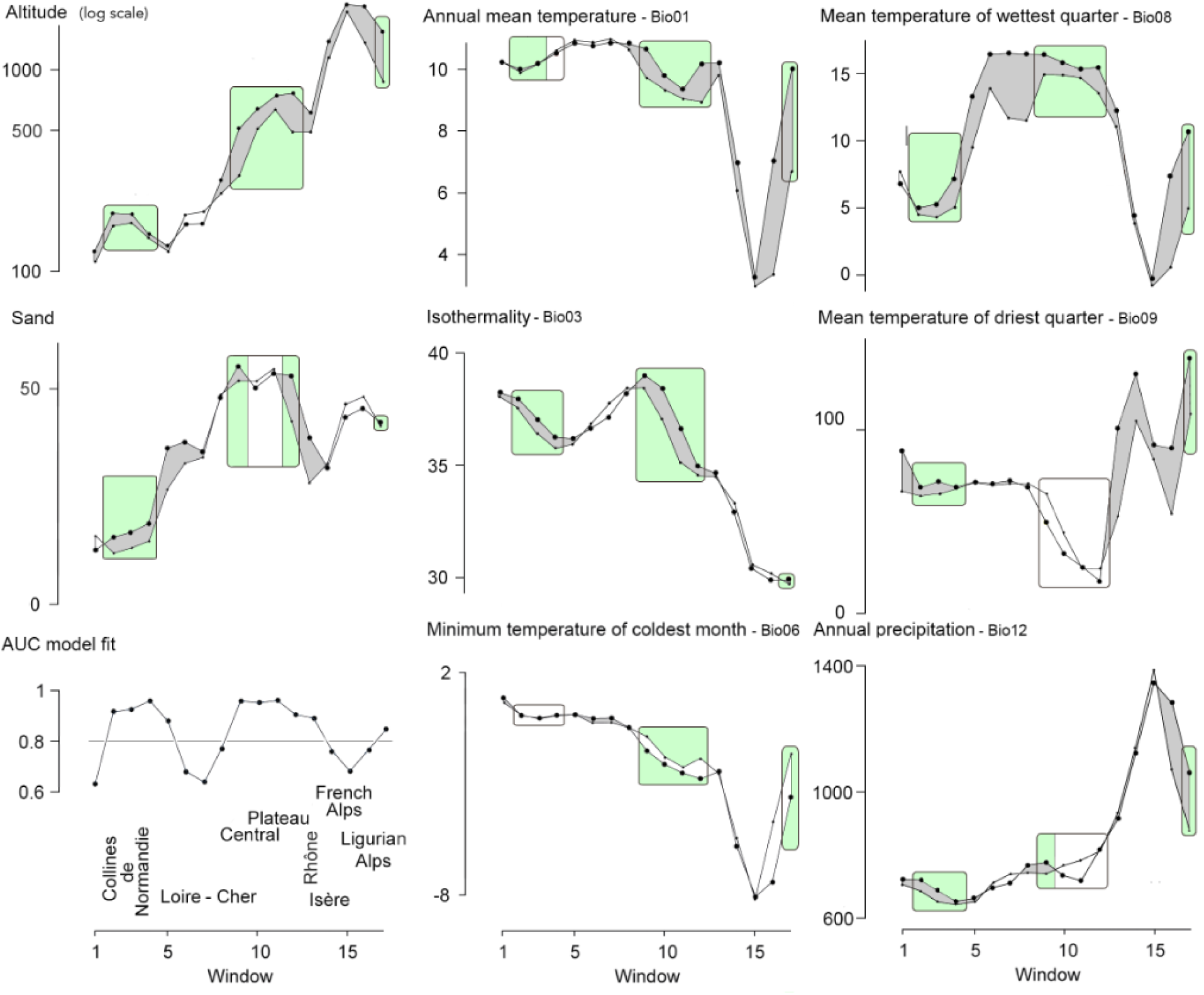
Average values for eight selected environmental variables over 17 windows that follow the *Bufo bufo – B. spinosus* hybrid zone from the Atlantic coast (window 1) to the Mediterranean (window 17). Units are as in table 1; see also Hijmans et al., 2005). Values for *B. bufo* and *B. spinosus* are shown by small and large dots, respectively. Grey areas indicate that values for *B. bufo* are lower than for *B. spinosus*. The graph bottom left provides AUC model fit values along with topographical references. Rectangles indicate stretches of the species contact for which the environmental models have good fit (AUC > 0.8), with consistent results indicated by green shadings. For the other windows with less than good model fit, signals are likely to be absent or void, either from poor sampling (window 1), the presence of rivers (windows 5-9, 13-14), or a thin or absent species’ contact (windows 16-17) (see figure 3).

## Discussion

We used two approaches to assess the role of environmental variables in shaping the common-spined toad hybrid zone. While both performed relatively well at predicting where pure populations will be, the global model performed poorly when genetically admixed populations were included. This may reflect the heterogeneity of the hybrid zone, where the spatial context of successive transects varies greatly in association with topography. Our results also highlight the sliding window approach as most useful for dealing with long, complex hybrid zones, because different variables will have different impact on hybrid zone structure in different sections of the zone, and these are highlighted as such.

Among selected environmental variables, most were related to temperature (bio01, bio03, bio06, bio08, bio09) or precipitation (bio12), but soil type also entered the global model. While the former may generally be associated with a differentiation between a northern (cooler, more humid) and a southern (warmer, drier and more extreme) species, the role of soil type is not obvious, and may have an indirect association with the distribution of the two toad species (e.g. through larval or postmetamorphic associations with pH and vegetal communities). However, land use categories did not make it into the models, but perhaps they operate at a smaller spatial scale and have little or no bearing on the wider position of the hybrid zone.

High model fit was also associated with higher altitudes for *B. spinosus* than for *B. bufo*. This observation is not compatible with the climate preferences alluded to in the above, with as a further complication that the critical altitude increases with an increasingly continental climate. Low model fit tended to be associated with less populated areas and with rivers. These are areas where dispersal is reduced and where hybrid zones are predicted to settle (Barton & Hewitt, 1985). The highest records for *B. bufo* are from the Swiss Alps at 2300 m (Malkmus & Grossenbacher, 2013) where the species is expanding its elevation range (Luscher et al., 2016). The highest records for *B. spinosus* are from the Pyrenees at 2600 m with, however, low population densities above 1500 m (Balcells, 1975; Ortiz Santaliestra, 2014). We were unable to sample from France to Italy and it is currently unclear to what degree the species are (or were) in contact, other than along the Mediterranean coast.

Our study shows that rivers can have a larger impact than climatic/environmental variables, suggesting that the presence of a river can break down environmental associations with toad presence. The low numerical model fit at rivers may simply reflect the fact that these linear structures were not incorporated in our two-dimensional modelling approach. Historical factors can also overrule the importance of environmental variables, for instance when different regions involve different intraspecific lineages, such as the northern Balkan lineage of *B. bufo* and its southern, Apennine counterpart (Garcia-Porta et al., 2012; Arntzen et al., 2017). The failure of the model to accurately predict the species present in Normandy and Brittany may be a ‘preoccupancy effects’, where a species is present in an area that is more environmentally favourable for the other species just because it has been long part of its range. Alternatively, *B. bufo* might have occupied these areas but did not make it because the colonization route was inaccessible.

Predicting the fate of moving hybrid zones is challenging because of the complex interaction between topography and environmental variables, but our sliding window approach, coupled with phylogeographic analyses, would seem to offer great potential to make predictions on species advances and retreats in a global change scenario.

## Acknowledgements

We thank Gaëlle Caublot, Jacob McAtear, Rob Veen and Annie Zuiderwijk for help with tissue collecting.

## References

Abbott, R., Albach, D., Ansell, S., Arntzen, J.W., Baird, S.J.E., Bierne, N., Boughman, J.W., Brelsford, A., Buerkle, C.A., Buggs, R., Butlin, R.K., Dieckmann, U., Eroukhmanoff, F., Grill, A., Cahan, S.H., Hermansen, J.S., Hewitt, G., Hudson, A.G., Jiggins, C., Jones, J., Keller, B., Marczewski, T., Mallet, J., Martinez-Rodriguez, P., Moest, M., Mullen, S., Nichols, R., Nolte, A.W., Parisod, C., Pfennig, K., Rice, A.M., Ritchie, M.G., Seifert, B., Smadja, C.M., Stelkens, R., Szymura, J.M., Vainola, R., Wolf J.B.W. & Zinner, D. (2013) Hybridization and speciation. Journal of Evolutionary Biology 26: 229–246.

Arntzen, J.W. (2019) An amphibian species pushed out of Britain by a moving hybrid zone. Manuscript under minor revision for Molecular Ecology.

Arntzen, J.W., de Vries, W., Canestrelli, D. & Martínez-Solano, I. (2017) Hybrid zone formation and contrasting outcomes of secondary contact over transects in common toads. Molecular Ecology 26: 5663–5675.

Arntzen, J.W., McAtear, J., Butôt, R. & Martínez-Solano, I. (2018) A common toad hybrid zone that runs from the Atlantic to the Mediterranean. Amphibia-Reptilia 39: 41–50.

Arntzen, J.W., Trujillo, T., Butôt, R., Vrieling, K. & Schaap, O.D., Gutiérrez-Rodriquez, J., Martinez-Solano, I. (2016) Concordant morphological and molecular clines in a contact zone of the common and spined toad (*Bufo bufo* and *B. spinosus*) in the northwest of France. Frontiers in Zoology 13: 1–12.

Balcells, E. (1975). Observaciones sobre el ciclo biológico de anfibios de alta montaña y su interés en la detección del inicio de la estación vegetativa. Publicaciones del Instituto de Biología Aplicada 7: 55–153.

Barton, N.H. & Hewitt, G.M. (1985) Analysis of hybrid zones. Annual Review of Ecology and Systematics 16: 113–148.

Fick, S.E. & R.J. Hijmans (2017) Worldclim 2: New 1-km spatial resolution climate surfaces for global land areas. International Journal of Climatology 37: 4302–4315.

Garcia-Porta, J., Litvinchuk, S.N., Crochet, P.A., Romano, A., Geniez, P.H., Lo-Valvo, M., Lymberakis, P. & Carranza, S. (2012) Molecular phylogenetics and historical biogeography of the west-palearctic common toads (*Bufo bufo* species complex). Molecular Phylogenetics and Evolution 63: 113–130.

Hijmans, R.J., Cameron, S.E., Parra, J.L., Jones, P.G. & Jarvis, A. (2005) Very high resolution interpolated climate surfaces for global land areas. International Journal of Climatology 25: 1965–1978.

Hunter, E.A., Matocq, M.D., Murphy, P.J. & Shoemaker, K.T. (2017) Differential effects of climate on survival rates drive hybrid zone movement. Current Biology 27: 3898–3903.

IBM SPSS (2016) Statistical Package for the Social Sciences. SPSS Inc., Chicago, USA.

ILWIS (2009). Integrated Land and Water Information System (ILWIS). Open software version 3.6. ITC, Enschede, The Netherlands.

Lescure, J. & de Massary, J.C. (2012) Atlas des Amphibiens et Reptiles de France. Biotope: Muséum National d’Histoire Naturelle. Paris, France.

Luscher, B., Beer, S. & Grossenbacher, K. (2016). Die Höhenverbreitung der Erdkrote (*Bufo bufo*) im Berner Oberland (Schweiz) unter sich verändernden Klimabedingungen. Zeitschrift für Feldherpetologie 23: 47–58.

Malkmus, R. & Grossenbacher, K. (2013) Fortpflanzungserfolg der Erdkröte (*Bufo bufo*) in hochalpinen Gewässern. Zeitschrift für Feldherpetologie 20: 102–104.

Mallet, J. (2005) Hybridization as an invasion of the genome. Trends in Ecology and Evolution 20: 229–237.

McQuillan, M.A., & Rice, A.M. (2015) Differential effects of climate and species interactions on range limits at a hybrid zone: potential direct and indirect impacts of climate change. Ecology and Evolution 5: 5120–5137.

Ortiz Santaliestra, M.E. (2014) Sapo común – *Bufo spinosus*. In: Enciclopedia Virtual de los Vertebrados Españoles. Salvador, A., Martínez-Solano, I. (Eds.). Museo Nacional de Ciencias Naturales, Madrid, Spain. http://www.vertebradosibericos.org/

Panagos P., Van Liedekerke M., Jones A. & Montanarella L. (2012) European Soil Data Centre: Response to European policy support and public data requirements. Land Use Policy 29: 329–338.

Pritchard, J.K., Stephens, M. & Donnelly, P. (2000) Inference of population structure using multilocus genotype data. Genetics 155: 945–959.

Recuero, E., Canestrelli, D., Vörös, J., Szabó, K., Poyarkov, N.A., Arntzen, J.W., Crnobrnja-Isailovic, J., Kidov, A.A., Cogălniceanu, D., Caputo, F.P. & Nascetti, G. (2012) Multilocus species tree analyses resolve the radiation of the widespread *Bufo bufo* species group (Anura, Bufonidae). Molecular Phylogenetics and Evolution 62: 71–86.

Riemsdijk, I. van, Arntzen, J.W., Butlin, R.K., Bucchiarelli, G., McCartney-Melstad, E., Rafajlovic, M., Scott, P., Toffelmier, E., Shaffer, B. & Wielstra, B. (2019a) Spatial variation in introgression along the common toad hybrid zone. Pp. 81–97, in I. van Riemsdijk, Hybrid Zone Dynamics in Amphibians. PhD-thesis. Naturalis Biodiversity Centre and the Institute of Biology at Leiden University, Leiden, The Netherlands.

Riemsdijk, I. van, Butlin, R.K., Wielstra, B. & Arntzen, J.W. (2019b) Testing an hypothesis of hybrid zone movement for toads in France. Molecular Ecology 28: 1070–1083.

Ryan, S.F., Deines, J.M., Scriber, J.M., Pfrender, M.E., Jones, S.E., Emrich, S.J. & Hellmann, J.J. (2018) Climate-mediated hybrid zone movement revealed with genomics, museum collection, and simulation modeling. Proceedings of the National Academy of Sciences 115: E2284–2291.

Systat (2007) Mystat 12. Systat Software, San Jose, USA.

Taylor, S.A., Larson, E.L. & Harrison. R.G. (2015) Hybrid zones: windows on climate change. Trends in Ecology and Evolution 30: 398–406.

